# Low-dose exposure to malathion and radiation culminates in the dysregulation of multiple neuronal processes instigating neurotoxicity and activation of neurodegeneration pathways in mice hippocampus

**DOI:** 10.1101/2023.06.08.544287

**Authors:** K N Rekha, B S Venkidesh, Sangeetha Nayak, Dinesh Reghunathan, Sandeep Mallya, Krishna Sharan, Rao B S Satish, Kamalesh Dattaram Mumbrekar

## Abstract

Neurodegenerative disorders are a debilitating and persistent threat to the global elderly population carrying grim outcomes. Their genesis is often multifactorial, with a history of early exposure to xenobiotics like pesticides or diagnostic exposure to ionizing radiation. A holistic molecular insight into their mechanistic induction is still unclear upon single or combinatorial exposure to different toxicants. In the present study, one-month-old C57/BL-6J male mice were treated orally with malathion (MAL) (50mg/kg body wt. for 14 days) and/or a single whole-body radiation (IR) (0.5 Gy) on the 8^th^ day. Post-treatment, behavioral assays were conducted to assess exploratory behavior, memory, and learning. Following sacrifice, brains were collected for histology, biochemical assays, and transcriptomic analysis. Differential expression analysis, Gene ontology, and pathway enrichment revealed several common and uniquely altered genes, biological processes, and pathways related to neurodegeneration, synaptic transmission and plasticity, neuronal survival, proliferation, and regulation of neuronal death. Increased astrogliosis was observed in the IR and co-exposure groups, with significant neuronal cell death and reduction in the expression of NeuN in all three groups. Sholl analysis and dendritic arborization/ spine density study revealed decreased total apical neuronal path length and dendritic spine density in all three groups. Decreased levels of antioxidant enzymes GST and GSH and acetylcholinesterase enzyme activity were also detected. However, there were no changes in exploratory behavior or learning and memory. Thus, explicating the molecular mechanisms behind MAL and IR can provide novel insights into the genesis of environmental factor-driven neurodegenerative pathogenesis.

## Introduction

Neurodegeneration constitutes the progressive deterioration in the morphology and function of neurons of selective brain regions that pathologically manifests in the form of various neurodegenerative disorders like Parkinson’s (PD), Alzheimer’s (AD), Amyotrophic lateral sclerosis (ALS), etc. Emerging studies suggest that the observed increase in neurodegenerative disorders over the years is proportional to the increased exposure to neurotoxic agents (Chin-Chan et al. 2015), with its impact varying in complexity, severity, and clinical significance. Xenobiotics, like various classes of drugs, cosmetics, pesticides, food additives, industrial chemicals, environmental pollutants, etc., are some common chemicals individuals can get exposed to in their lifetime (Patterson et al. 2010). Among these etiological factors, xenobiotics like pesticides and ionizing radiation are at the forefront, especially due to their concerning and increased application (Walther et al., 2023).

MAL is a broad-spectrum organophosphate used mainly for agricultural, domestic, and public health purposes, and owing to its slightly less toxic nature, it is extensively and sometimes recklessly overused in several parts of the world (Badr 2020). In non-target organisms, the main mechanism of action of MAL is through the inhibition of AchE activity in the brain, leading to adverse effects like a decrease in the strength of cranial motor nerve, proximal limb muscle, neck flexors, restlessness, hyperexcitability, seizure and in extreme cases, death (Tchounwou et al. 2015). Apart from this, MAL has been known to generate free radicals and cause antioxidative damage (Fortunato et al. 2006) and DNA damage (Réus et al. 2008) in rodent brains. MAL also increases pro-inflammatory cytokines like TNF-α (Tumor necrosis factor α) and IL-6 (Interleukin 6), which mediates neuronal damage (Mohammadzadeh et al. 2018), resulting in cognitive impairment and causes neuroinflammation by the activation of microglia leading to neuronal death (Ahmed et al. 2017). Further, it interferes with axonal transport via the alteration of motor proteins (Naughton and Terry 2018), induces mitochondrial dysfunction by altering the levels of mitochondrial complex activities, generation of ROS, ATP depletion, DNA fragmentation, and apoptosis (Venkatesan et al. 2017) in rodents and neuronal cell lines.

Medical usage of radiation contributes about 98% to the total population dosage through artificial sources while representing 20% of the total population exposure (WHO, 2022). Further, it has been studied that a single fraction of head and neck cancer radiotherapy can contribute up to 154.8 mGy to the out-of-field regions of the hippocampus (Auerbach et al., 2023).

Low-dose IR is implicated in inducing impairment in spontaneous behavior, habituation capacity, spatial memory, and logical reasoning and also causes learning deficits (Kiffer et al., 2020). It also causes morphological changes like a reduction in the complexity of the dendritic architecture and dendritic spine density (Parihar et al. 2015). Studies have also shown that it induces neurotoxicity by a decline in the hippocampal neurogenesis and neuroinflammation through increased pro-inflammatory markers like Iba-1, Cd68, and Cd11c (Acharya et al. 2015). Further, it alters oxidative phosphorylation, reduces mitochondrial complex enzyme and antioxidant levels, and causes mitochondrial dysfunction (Casciati et al. 2016). Expression of genes playing an important role in neuronal plasticity and signal transduction is also reduced on low-dose IR exposure (Casciati et al. 2016; Kempf et al. 2015).

There is a huge lack of understanding on how complex xenobiotics interact in combined exposure setting and if low-dose xenobiotics, which are generally less adverse, can have a large detrimental impact on human health when combined with other chemicals (Silins and Högberg 2011a). Both MAL and low-dose IR have been known to cause DNA damage through the generation of free radicals and decreased levels of antioxidant enzymes leading to apoptosis, with several reports showing that they induce neurotoxicity. Moreover, the LDR exposure effect often remains unpredictable and ambiguous due to various factors like dose and duration of exposure and inter-individual differences, etc. (Narasimhamurthy et al. 2022). Also, though MAL is predicted to play a role in the etiology of neurodegenerative diseases, its non-cholinergic mechanism at the level of pathways is still elusive, and several of the mechanistic studies conducted so far are majorly in *in-vitro* models. The present study elucidates the mechanism behind pesticide/IR exposure-induced neurotoxicity by employing high throughput methods using a mouse model. We also investigate if exposure to these agents at a young age can drive the organism toward neurodegenerative outcomes. Further, we also seek to understand how these two commonly acting xenobiotics may act in cases of combined exposure which is especially relevant when applied to a large population.

## Material and Methods

### Animals and treatment conditions

The animal model used for the current study was C57/BL male mice and was obtained from an in-house Central Animal Research Facility, Kasturba Medical College, Manipal Academy of Higher Education, Manipal. The ethical clearance was duly obtained from the Institutional Animal Ethics Committee (IAEC/KMC/108/2019 dated 11.09.2019). The animals were maintained in polypropylene cages under optimum temperature (20°C ± 2 °C) and light conditions (12h light and dark cycle). Sterile food and water were provided *ad libitum*.

4–5-week-old animals weighing 25-30g were used for the experiment, and body weight was monitored daily throughout the treatment. Thirty-six animals were divided into four groups containing nine animals each, namely; Control, MAL (50 mg/kg), Radiation (0.5 Gy), and Co-exposure (50 mg/kg MAL orally + 0.5 Gy Radiation). Commercially procured MAL (MP Biomedicals, India) was administered orally in saline through gavage for 14 days continuously, and on the 8^th^ day, a single whole-body exposure to 0.5 Gy of X-ray radiation using Versa HD Linear Accelerator delivering 6MV Photon Energy (Elekta, Sweden) was given. After the completion of treatment, the animals were subjected to behavioral assays followed by tissue collection for histological, biochemical, and molecular analysis.

### Sholl analysis and dendritic arborization by Golgi-Cox Staining

As suggested previously, the procedure was carried out (Narayanan et al. 2014). After Golgi-Cox staining, the brain was embedded in 2% low melting agarose, and sections of 150 µm thickness were taken using a vibratome (company, country), and images were captured using Olympus CKX53 (Olympus, Japan) microscope. Neurons were traced using the Simple Neurite Tracer plugin in Fiji. Sholl analysis was then performed on the traced neuron using the Sholl analysis plugin. The total number of average intersections, the number of intersections at different distances from the soma, as well as the total path length in both apical and basal neurons were calculated. Further, the spine density in every 10 µm of dendrite was also calculated.

### Nissl Staining for neuronal survival

Nissl staining was done according to the protocol described previously (Venkidesh et al., 2023). The images were captured using Olympus CKX53 (Olympus, Japan) microscope, and the image analysis was done using the Fiji plugin Cell counter to score the pyknotic cells. Three animals per group, with three sections per each animal, were analysed.

### Immunohistochemistry for mature neurons and astroglia activation

Immunohistochemistry was performed as previously described (Venkidesh et al. 2023) using NeuN (1:500 dilution, Invitrogen, USA) and GFAP (1:2000 dilution, Invitrogen, USA) markers for mature neurons and astroglia, respectively and with secondary antibody (1:1000 dilution, goat anti-rabbit) (Jackson ImmunoReasearch Laboratory, USA). Three animals per group, with three sections per each animal, were analysed. From each region, a minimum of three images were captured for the analysis, and from each section; a minimum of six images were captured on the whole. The images were captured using Olympus CKX53 (Japan), and the image analysis was done using the Fiji and Fiji plugin Cell counter. Further, the expression of NeuN was represented as % positively expressed area (Insausti et al. 2015).

### AChE inhibition assay

Ellman’s assay, with some modifications, was conducted to assess the levels of acetylcholinesterase inhibition (Banasik et al. 2003). The absorbance/ min was calculated using the slope, and the rate of the reaction was calculated using the formula;

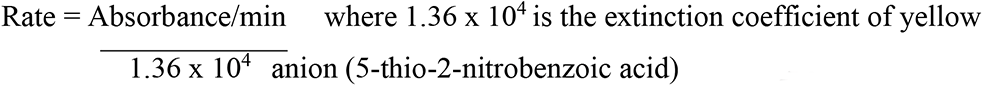

Further, the treatment groups were normalized with control and expressed as percentage normalized activity with respect to control.

### GSH and GST assay

The intracellular levels of GSH and GST were determined by protocols as described before (Das et al. 2016). The GSH and GST values were normalized with the respective protein values measured through Bradford assay and expressed in terms of μM/ mg protein.

### Open Field Test for anxiety-related behavior

The animal was placed in the center of an open square field facing the side away from the experimenter and allowed to explore for 5 minutes which was video recorded. After 5 minutes, the animal was removed, and the field was cleaned using 70% alcohol and allowed to dry. The video was then blind-scored for the duration spent in the center (Seibenhener & Wooten, 2015).

### Novel Object Recognition Test for recognition memory

The test was performed as described previously (Leger et al. 2013), and the experiment was recorded. The video was then blindly scored, and the parameters like duration and frequency of exploration of the novel as well as the familiar object, were noted. The discrimination index and recognition index were calculated from the data obtained.

### mRNA isolation, cDNA conversion, library preparation, and sequencing

The hippocampus was isolated from freshly dissected mice brains on ice, weighed, and then homogenized with TRIzol (Ambion, USA). The RNA was isolated according to the manufacturer’s protocol, and quality was checked by running on a gel. mRNA isolation and purification were performed using the Dynabead mRNA DIRECT Purification Kit (Invitrogen, USA), cDNA conversion, amplification, and library preparation were done using Ion Total RNA-Seq Kit v2 (Invitrogen, USA) according to the manufacturer’s protocol. The template-positive ion sphere particles (ISPs) were generated using the Ion PI Hi-Q OT2 200 Kit (Thermo Fischer, USA). The sequencing was carried out using 200bp sequencing chemistry in an Ion Proton semiconductor sequencing using an Ion PI chip manufacturer instruction (Invitrogen, USA). Two samples from each group were separately analyzed with no sample pooling. After the run, the transcriptomic data was downloaded in the form of ubam format and subjected to further bioinformatic analysis.

### Bioinformatic analysis

The fastq files obtained following the transcriptomic run of the mice hippocampal tissue were analyzed using nf-core/RNAseq pipeline version 3.10.1 on Nextflow version 22.10.4 1–3. The quality of the raw fastq files was assessed using FastQC and reported using MultiQC. The reads were mapped to the mouse genome (GRCm38) using STAR aligner, and the counts were extracted using featureCounts software4–6. Differential expression (DE) analysis between groups was performed using EdgeR, an R Bioconductor package 7. A p-value of <=0.05 and fold change of >= 1.5 and <= - 1.5 was defined to establish significant upregulated and downregulated genes, respectively.

The significant differentially expressed genes (DEGs) were then subjected to gene ontology enrichment analysis and pathway enrichment analysis using the online modules at SR plot (https://www.bioinformatics.com.cn/en) through the Cluster profiler (Yu et al. 2012) and Pathview packages (Luo and Brouwer 2013). A STRING network (Szklarczyk et al. 2023) to visualize the functional interaction of the DEGs in each treatment group was constructed at a high confidence score (0.7), and only the significant and connected nodes within our DEG network were visualized. The network was imported to Cytoscape (Shannon et al. 2003) and was modified using Cytohubba (Chin et al. 2014) to represent the network interaction of the top 50 genes.

### Statistical analysis

Statistical analysis was carried out using the GraphPad Prism v.8 software, and one-way ANOVA was used to analyze the data. Data were expressed in terms of mean ± SD or mean ± SEM, and a p-value less than 0.05 was considered significant.

## Results

### Neuronal death

Neuronal death was assessed, and the number of pyknotic cells in the hippocampal region was assessed (Fig 1 a-d). Pyknotic cells were found to be significantly increased in 0.5Gy (p>0.001) and MAL (p>0.05) treated group as well as the in the co-exposure group with the highest number of pyknotic cells in the co-exposure group (p>0.0001) (Fig d).

**Fig 1:**
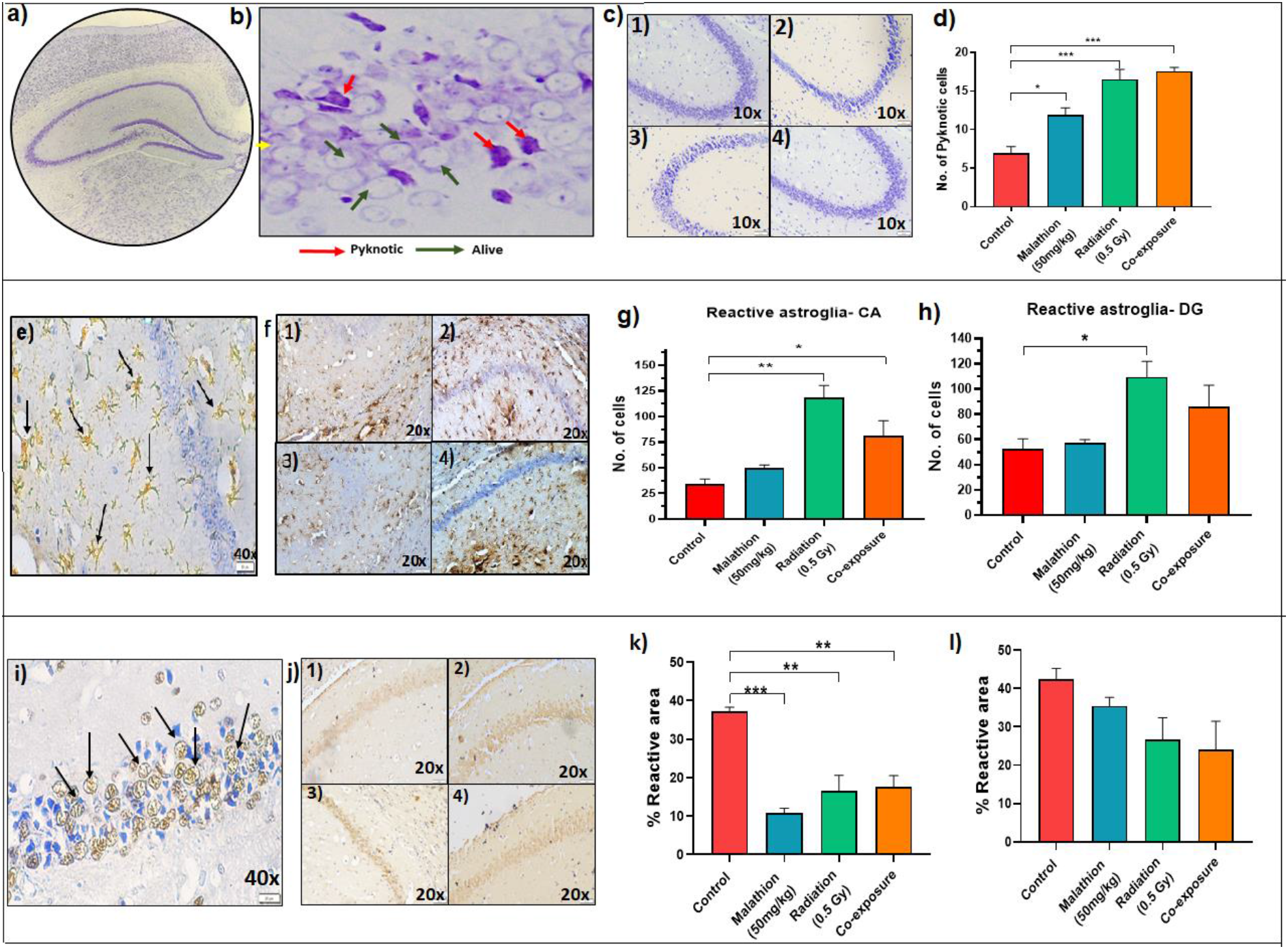
Effect of radiation and malathion on neuronal survival, inflammation, neuronal maturation, enzyme activity and behavior. *a) Nissl-stained mice hippocampus; b) representative image showing pyknotic and live neurons in the mice hippocampus; c) representative images of hippocampal regions of different groups - 1) control 2) radiation 3) malathion 4) co-exposure; d) graph depicting the number of pyknotic neurons in different groups (n=3, p≥*0.05, **0.001, data represented as ± SEM). e) Representative image showing reactive astroglia; f) representative images of GFAP positive cells in the hippocampus of different groups 1) control 2) radiation 3) malathion 4) co-exposure; g) graph depicting the number of reactive astroglia in CA region; h) graph representing the number of reactive astroglia in DG region (n=3, p≥*0.05, **0.01, data represented as ± SEM). i) Representative image showing NeuN positive cells; j) representative images of NeuN positive cells in the hippocampus of different groups 1) control 2) radiation 3) malathion 4) co-exposure; k) graph depicting the percentage of NeuN positive area in the DG region; l) graph representing the percentage of NeuN positive area in CA region (n=3, p≥*0.05, **0.01, ***0.001, data represented as ± SEM).*

### Astroglia activation and neuronal maturation

In the cornu ammonis (CA), the expression of reactive GFAP increased in 0.5Gy (p>0.01) and co-exposure groups (p>0.05) (Fig 1g). In the dentate gyrus (DG) region, increased astrogliosis was observed in the 0.5Gy group (p>0.05) (Fig 1h). Further, the glial cell projections were higher in number as well as increased in their thickness in the treatment groups (Fig 1 e,f). NeuN expression was also calculated in the DG and CA region (Fig 1 i and j). NeuN expression decreased in MAL (p>0.001), 0.5Gy (p>0.01), and co-exposure groups (p>0.01) (Fig 1k) in the DG region, indicating a decreased number of mature neurons in the hippocampal region.

### Dendritic arborization and spine density

Compared to the control, apical dendritic path length showed a decreasing trend in 0.5Gy (p> 0.05), MAL (p>0.05), and co-exposure (p>0.01) (Fig 2e). No significant changes were observed in the total basal dendritic path length and the average number of apical and basal dendritic branches (Fig 2 f-g). Further, though there were no changes in the number of basal dendritic intersections (Fig 2i), a decrease in the third intersection in the 0.5 Gy and MAL group, 4^th^ intersection in the co-exposure group and 6^th^ intersection in the 0.5 Gy and co-exposure groups compared to control was observed in the number of apical intersections (Fig 2j) (p> 0.05). Further, both 0.5Gy and co-exposure groups showed a decrease in the dendritic spine density (Fig 2k-m) (p> 0.05).

**Fig 2:**
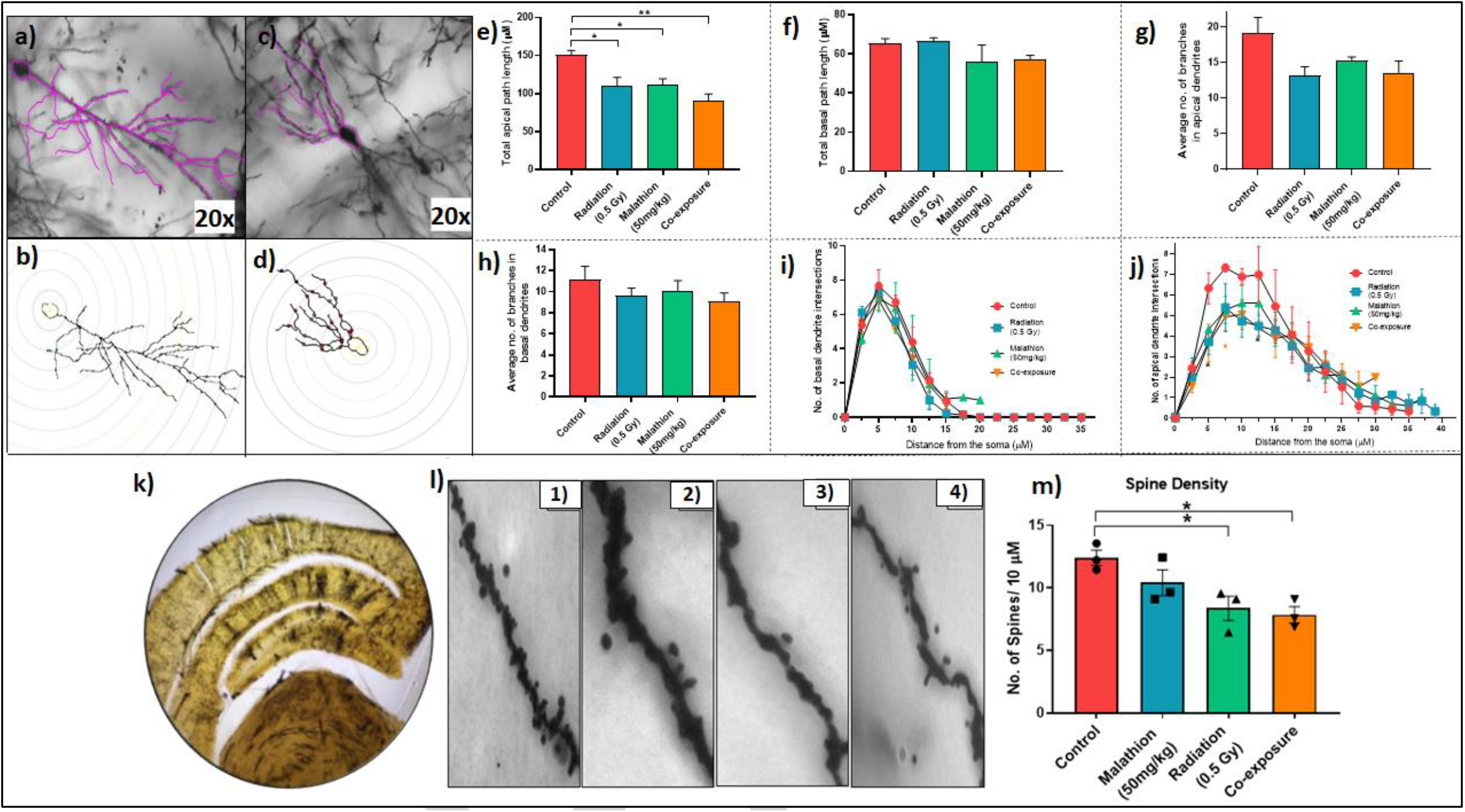
Effect of radiation and malathion on neuronal morphology and arborization. *a) Representative image of traced apical dendrite of a pyramidal neuron; b) Representative image of apical dendritic neuron after undergoing Sholl analysis; c) Representative image of traced basal dendrite of a pyramidal neuron; d) Representative image of basal dendritic neuron after undergoing Sholl analysis; e) Graph depicting total apical dendritic path length; f) Graph depicting total basal dendritic path length; g) Graph depicting the average number of branches in the apical dendrites; h) Graph depicting the average number of branches in the basal dendrites; i) Graph depicting number of intersections in the basal dendrites; j) Graph depicting number of intersection in the apical dendrites; k) Golgi-cox stained mice hippocampus; l) Representative images of dendritic spines in – 1) Control, 2) Malathion, 3) Radiation, 4) Co-exposure groups; m) Graph depicting the change in spine density between different groups (n=3, p≥*0.05, **0.01, ***0.001, data represented as ± SEM)*.

### Antioxidant capacity

GSH levels were significantly lowered in the 0.5Gy and co-exposure group when compared to the control (p>0.05) (Fig 3a). Similarly, GST levels were also significantly lowered in all three treatment groups with respect to control (p> 0.001) (Fig 3b).

**Fig 3:**
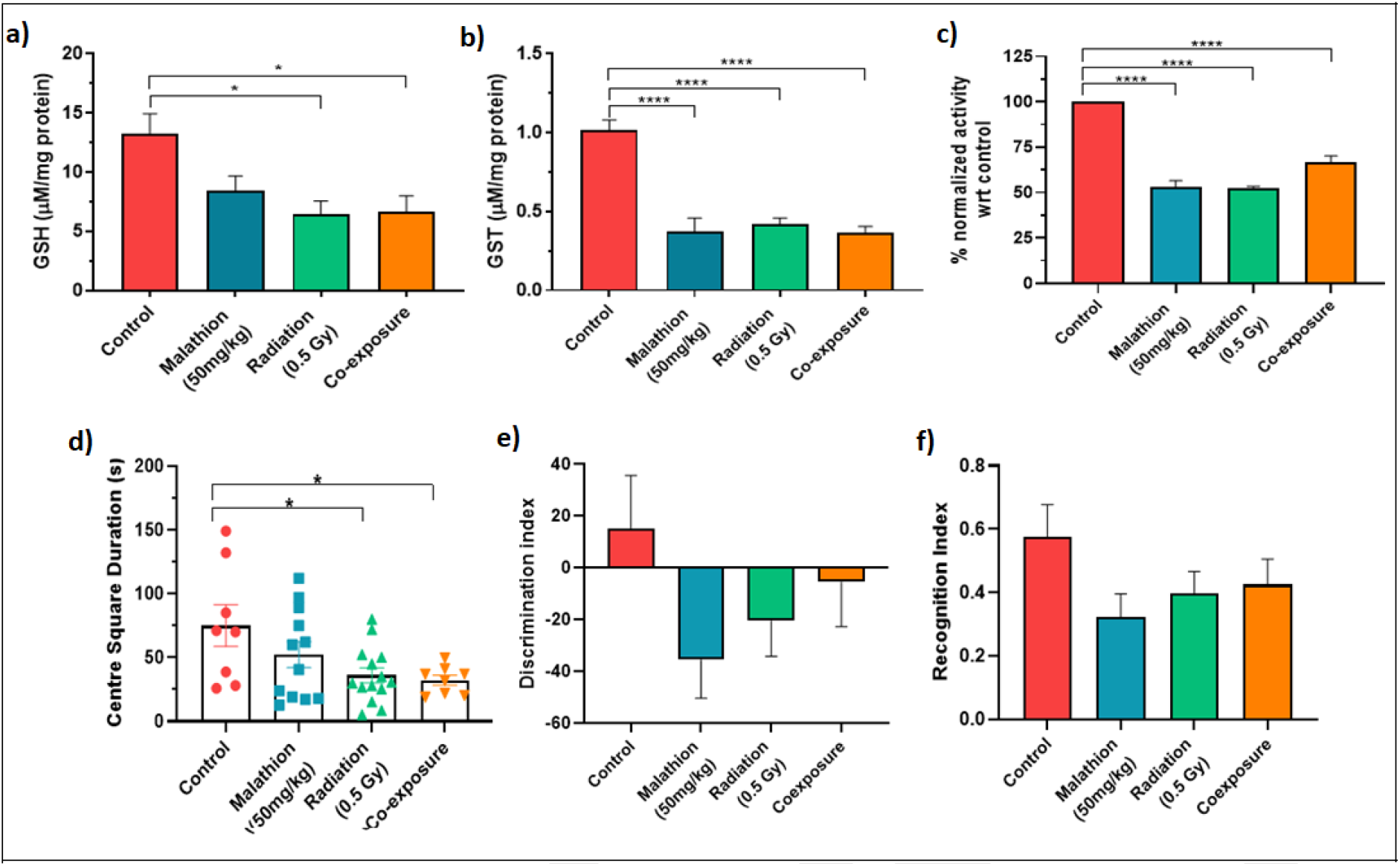
Effect of radiation and malathion on enzyme activity and behavior. *Graphs depicting a) GSH enzyme levels between groups; b) GST enzyme levels between groups; c) % acetylcholinesterase enzyme inhibition activity between groups (n=6, p≥*0.05, ****0.0001, data represented as ± SEM). Graphs depicting; d) center square duration in open field test; e) determination index in novel object recognition; f) recognition index in novel object recognition (n=9)*.

### AChE inhibition

The levels of acetylcholinesterase were significantly reduced in all three treatment groups (p> 0.001) (Fig 3c).

### Behavioral changes

All three treatment groups, 0.5Gy, MAL, and co-exposure, did not show any significant changes in exploratory behavior (Fig 3d). There was no significant change observed in both the discrimination and recognition index upon the introduction of the novel object to the box in any of the treatment groups (Fig 3e and 3f).

### Transcriptomic analysis

Differential expression analysis revealed 18,967 differentially expressed genes (DEGs) with respect to control, out of which, in the 0.5Gy group, 246 genes were significantly downregulated, and 157 genes were significantly upregulated (Fig 4a). In the MAL treatment group, 339 genes were downregulated, and 487 genes were upregulated significantly (Fig 4b), while in the co-exposure group, 322 genes were downregulated and 294 genes were upregulated significantly (Fig 4c).

**Fig 4:**
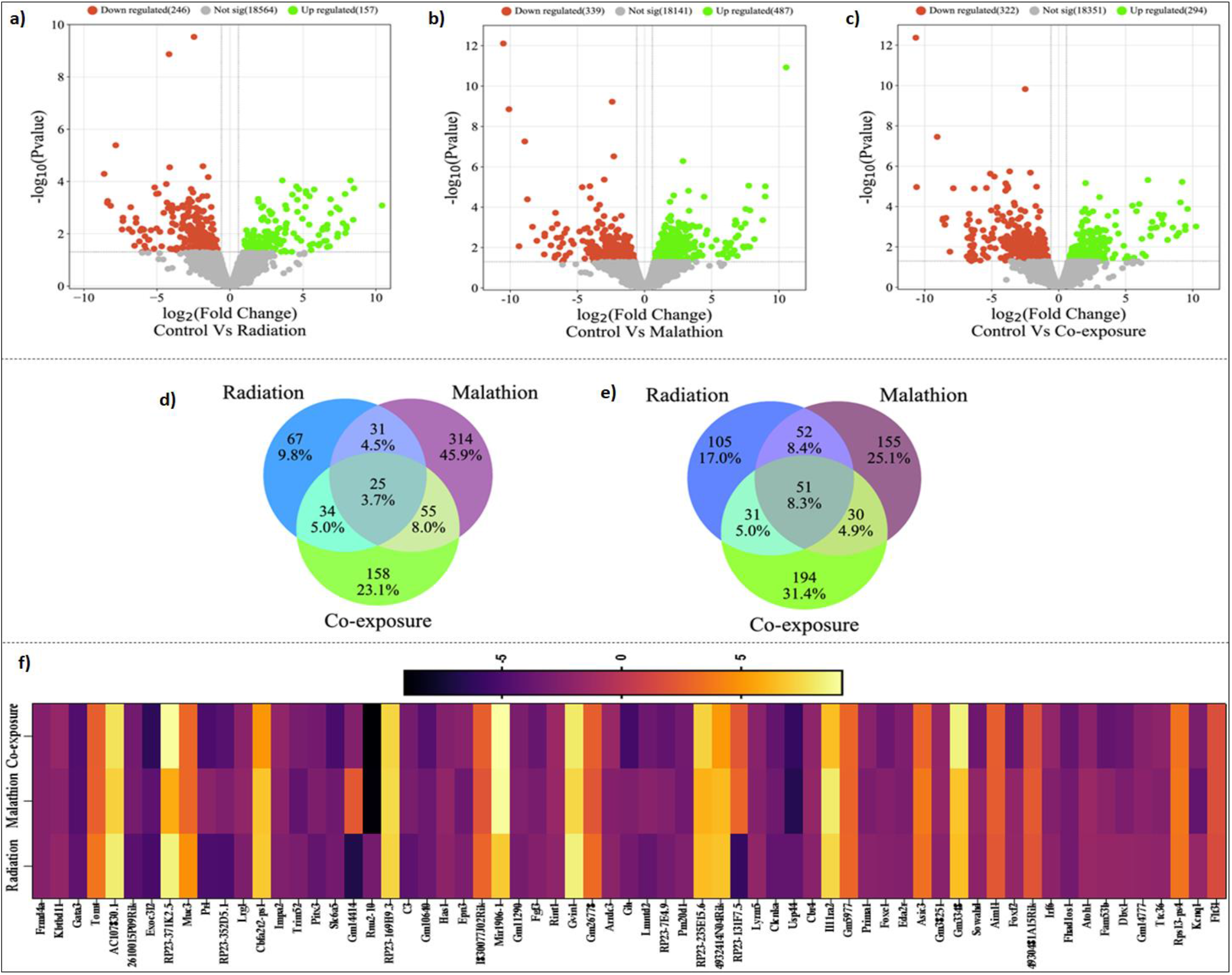
Differentially expressed genes post exposure to radiation and malathion. *Volcano plot showing the number of significant DEGs in - a) Radiation, b) Malathion, c) Co-exposure groups. Venn diagram displaying the number of DEGs in all three groups that are- d) Upregulated, e) Downregulated; f) Heat map depicting the number of commonly dysregulated genes in all the three groups and their respective expression pattern expressed in terms of fold change value (n=2, p≥0.05 values considered significant)*

25 genes were commonly upregulated between all three treatment groups (Fig 4d, e), and 51 genes were commonly downregulated between all three treatment groups (Fig 4d, e). After filtering the genes according to the threshold, 65 genes in total were found to be common, and these DEGs and their respective fold change variation between the three different groups were represented in a heat map (Fig 4f).

#### Gene ontology and pathway and STRING enrichment analysis

GO enrichment revealed that 262 common BPs, 15 common CCs, and 29 common MFs were affected between 0.5Gy, MAL, and co-exposure groups. 303 BPs, 45 CCs, and 44 MFs were common between 0.5Gy and the MAL group (Fig 5 a-c). The functionally enriched protein-protein interaction network for the altered DEGs is depicted in Fig 7. The Cnet plot for top 10 enriched biological processes and pathways can be found in the supplementary figure file.

**Fig 5:**
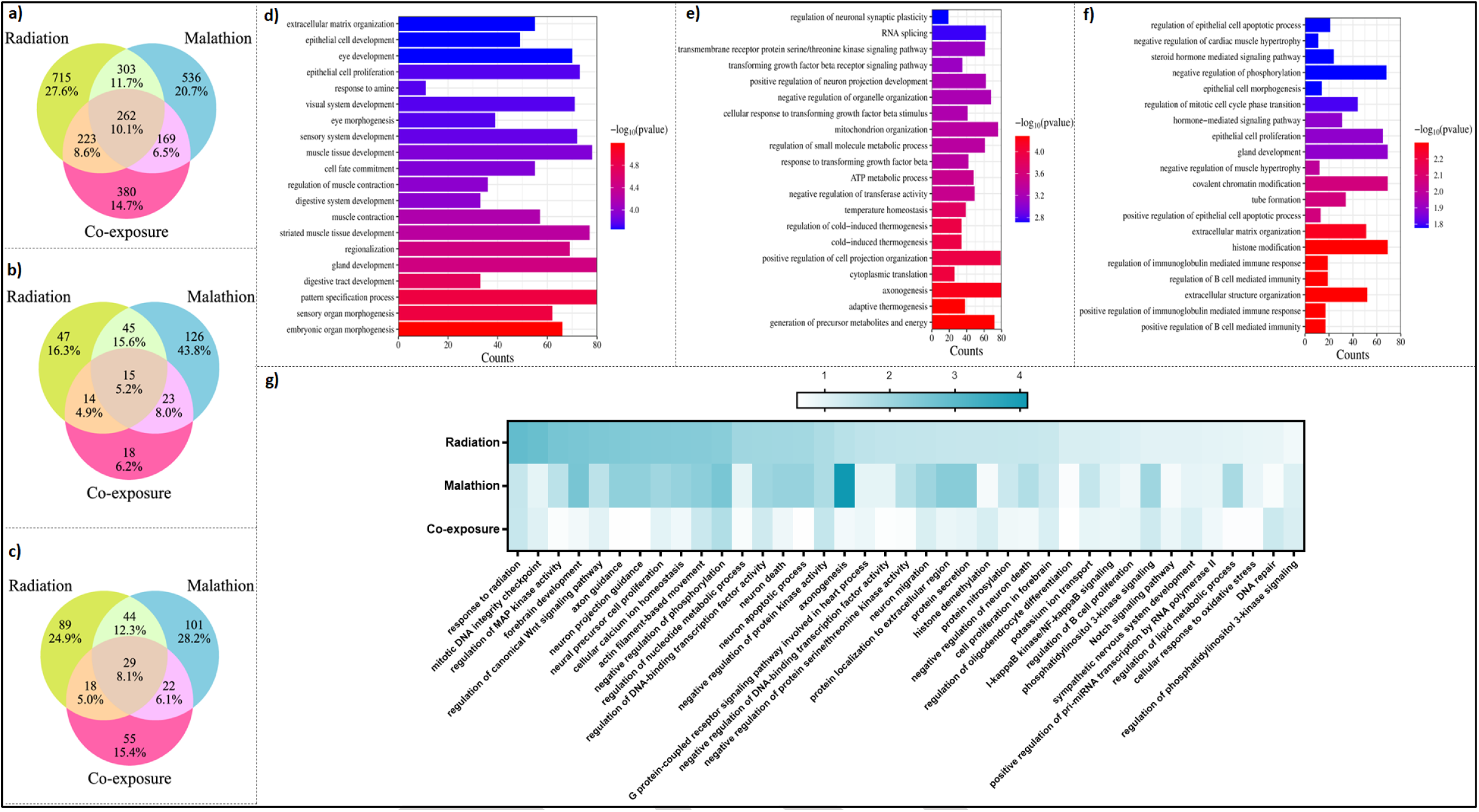
Biological processes altered post exposure to radiation and malathion. *Venn diagram displaying the number of a) BPs altered in all the three groups, b) CCs altered in all the three groups, c) MFs altered in all the three groups. Top 20 enriched biological processes affected in d) Radiation, e) Malathion, f) Co-exposure. g) Selected common processes in all the three groups and their enrichment value represented as a heat map (n=2, p≥0.05 values considered significant)*

##### Radiation single exposure

Gene ontology (GO) enrichment showed that 1503 significant biological processes, 121 cellular components, and 180 molecular functions were affected. The top twenty biological processes affected are plotted in a bar graph with their enrichment value and gene counts. 223 BPs, 14 CCs, and 18 CCs were common between 0.5Gy alone and the co-exposure group (Fig 5 a-c). The top 20 altered biological processes (Fig 5d) include processes like extracellular matrix organization, epithelial cell development and proliferation, response to amine, sensory system development, etc.

73 pathways were significantly affected in the 0.5Gy group. Some of the major pathways like the MAPK, PI3K-Akt, Apelin, NF-κB, TGF-β, cAMP and Notch signaling pathway, Cholinergic synapse, Serotonergic synapse, Dopaminergic synapse Axon guidance, etc., which play an important role in regulating neuronal functioning and survival were altered (Fig 6a). The STRING PPI of the 0.5Gy group had 225 nodes, 32 edges with a PPI enrichment value of 0.00332 (Fig 7a).

**Fig 6:**
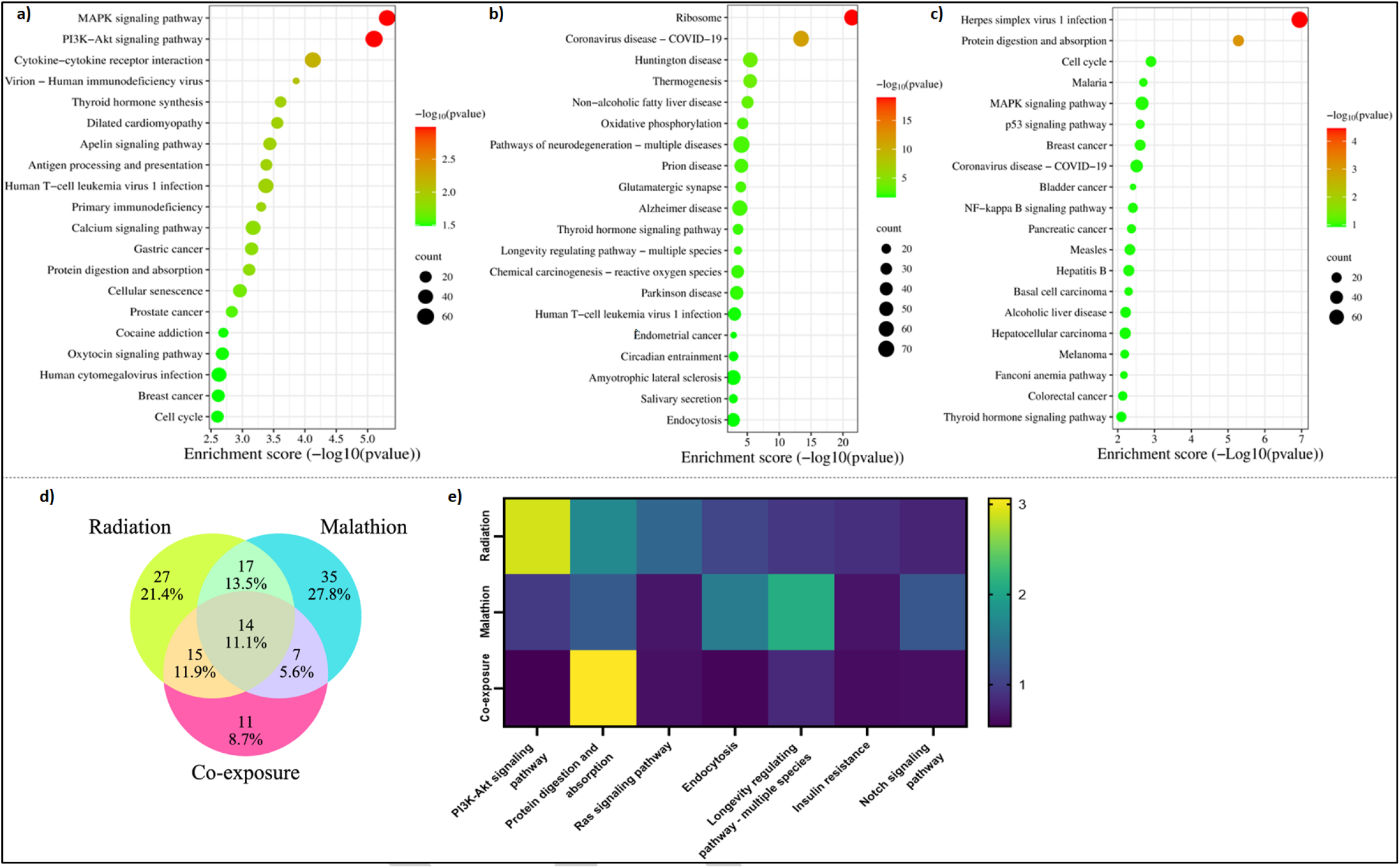
Pathways altered post exposure to radiation and malathion. *Top 20 enriched pathways affected in a) Radiation, b) Malathion, c) Co-exposure. d) Venn diagram displaying the number common and unique pathways altered in all the three groups. g) Selected common pathways altered in all the three groups and their respective enrichment value represented as a heat map (n=2, p≥0.05 values considered significant)*

**Fig 7:**
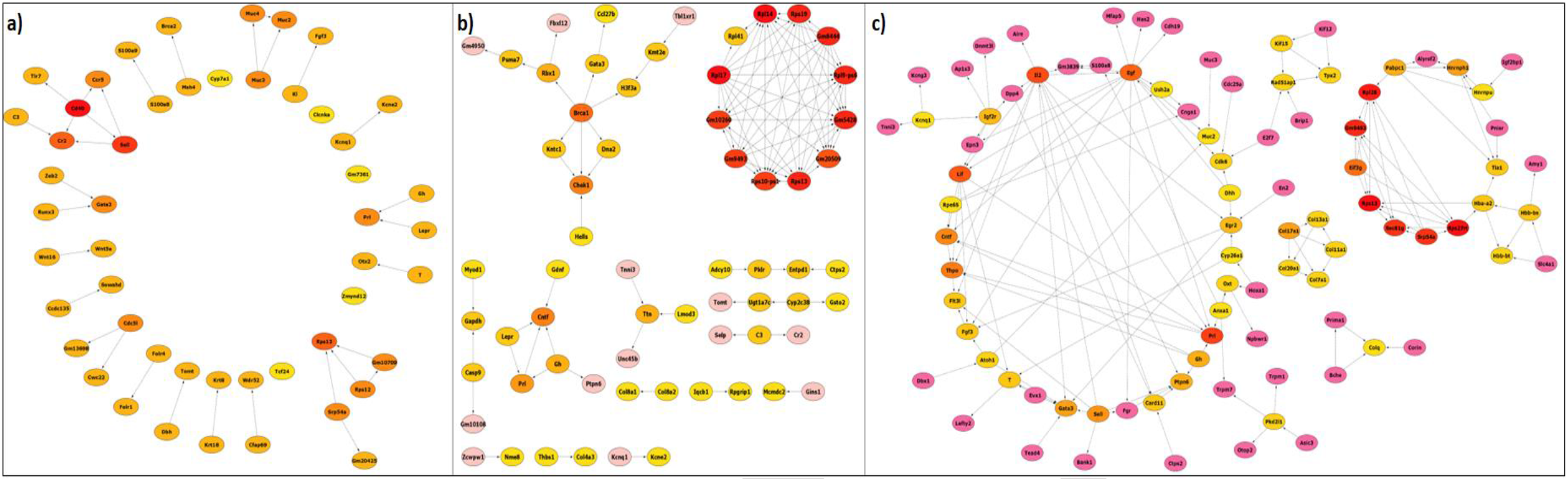
STRING network depicting PPI interactions between genes post exposure to radiation and malathion. *Functional protein-protein interaction network of the DGEs of a) Radiation, b) Malathion, c) Co-exposure groups, a high confidence score of 0.7 was considered, the color intensity of the nodes represents the confidence of the interaction and the arrows connecting the nodes detect the targets, closely associated and interacting nodes are connected by lines (n=2, p≥0.05 values considered significant)*.

##### MAL single exposure

GO enrichment revealed that 1270 significant biological processes, 209 cellular components, and 196 molecular functions were affected. The top twenty biological processes affected are plotted in a bar graph with their enrichment value and gene counts. 223 BPs, 23 CCs, and 22 MFs were common between MAL and co-exposure groups. The top 20 altered biological processes included regulation of neuronal synaptic plasticity, RNA splicing, mitochondrion organization, axonogenesis, etc. (Fig 5e).

73 pathways were significantly affected in the 0.5Gy group. Some of the major pathways like Ribosome, Huntington’s disease, Oxidative phosphorylation, Pathways of neurodegeneration - multiple diseases, Glutamatergic synapse, Alzheimer’s disease, Parkinson’s disease, Amyotrophic lateral sclerosis, Notch signaling pathway, Axon guidance, etc., which play an important role in neurodegenerative pathogenesis were altered (Fig 6b). The STRING PPI network of MAL had 446 nodes and 676 edges with a PPI enrichment value of 1^-16^ (Fig 7b).

##### Control vs. Co-exposure

GO enrichment revealed 1034 significant biological processes, 70 cellular components, and 124 molecular functions were affected. 380 BPs, 18 CCs, and 55 MFs affected were unique to the co-exposure group. The top 20 altered biological processes included regulation of epithelial cell apoptotic process, regulation of immunoglobin-mediated immune response, histone modification, etc. (Fig 5f).

47 pathways were significantly affected in the 0.5Gy group. Some of the major signaling pathways like MAPK, p53, NF-κB, JAK-STAT, Notch, Rap1, Hippo, FoxO, PI3K-Akt, etc., which play an important role in immune response, neuronal proliferation and survival, and synaptic plasticity were altered (Fig 6c). The STING PPI network of the co-exposure group had 341 nodes and 97 edges with a PPI enrichment value of 0.000558 (Fig 7c).

## Discussion and conclusion

The role of radiation and pesticides in neurodegenerative disorders and neurotoxic symptoms is well known. However, there is limited information available on the molecular mechanism behind the effect of low-dose radiation or pesticides and how early exposure can contribute to neurotoxicity. The present study aimed to elucidate the mechanism involved in the manifestation of neurotoxic and the subsequent neurodegenerative effects post-exposure to IR and MAL (Lowe et al. 2009; Vellingiri et al. 2022). In the present study, inhibition in acetylcholinesterase enzyme activity was seen in the 0.5Gy and MAL treatment groups. Previously MAL has been linked to AChE inhibition (Trevisan et al. 2008); however, studies looking at the effect of radiation on the same are sparse.

One of the earliest cellular responses to neurotoxic stimuli involves the initiation of neuroinflammation, typically through the activation of resident microglia and astrocytes, which constantly monitor the brain microenvironment for the presence of damaged or dead cells, contribute to synaptic plasticity, and maintain homeostasis (Gelders et al. 2018). While in IR and co-exposure, reactive astrogliosis was increased, MAL treatment did not cause any changes to the astroglia, suggesting that at the current dose, some other response, likely involving oxidative stress response machinery, may have contributed to the neurotoxicity associated with MAL exposure (Réus et al. 2008). Though MAL alone did not cause any astrogliosis, co-exposure showed an additive effect. We further noted a significant decrease in GSH in IR and co-exposure groups and reduced levels of GST in all three treatment groups. Similar results were seen with MAL administration in rats (Trevisan et al. 2008). The involvement of redox machinery and the detoxification of free radicals by the antioxidant enzymes have been previously linked to neurodegeneration (Nakamura et al. 2012), indicating that single or combined exposure to MAL and IR can induce similar effects.

Neuronal viability in the brain is closely associated with conditions involving tissue injury or neurodegeneration (Morrison et al. 2002) and is also one of the key hallmarks of neurodegenerative diseases (Hirsch 1999). In our study, MAL treatment hampered neuronal survival, as indicated by an increase in the pyknotic neurons. Likewise, low-dose IR has been associated with neuronal cell death (Kovalchuk and Kolb 2017), and in the present study, the pyknotic cells were significantly higher in both IR and co-exposure groups. Further, we also looked at the percentage expression of surviving mature neurons using the NeUN marker and found that the expression of surviving neurons was also reduced in the MAL, IR, and co-exposure group significantly in the DG region. An interesting thing to note is that out of all the three treatment groups, the co-exposure group showed the highest number of pyknotic cells and the lowest number of viable neurons, which indicates that the combined treatment of MAL and IR produced an overall synergistic effect on the neurons. Similar phenomena with both synergistic and antagonistic effects have been shown to occur whenever environmental chemicals and other stressors have been known to combine (Holmstrup et al., 2010). Varied interaction may be seen when individual chemicals with similar or dissimilar modes of action may affect the toxicity of each other either through synergism or dose/ effect addition or antagonism depending on the dose and time window of the interaction (Silins and Högberg 2011b) or show completely different effect than observed singularly (Arif et al., 1994). In our case, though the astroglia activation wasn’t the highest compared to IR, the antioxidant damage and cell death showed synergistic damage, while the percentage of acetylcholinesterase enzyme inhibition activity with respect to control was at closely comparable levels of MAL and IR groups showing a slight antagonistic effect. This just affirms that the effect of the interplay of various neurotoxicants is contingent upon the dose, duration, and physiological variations within the exposed individual, making it difficult to predict the response. Therefore, increased oxidative stress, increased neuroinflammation, decreased viability, and increased pyknotic cells may pose a devastating effect on neuronal homeostasis.

Dendrites and dendritic spines are crucial in synaptic transmission, controlling behavior and memory, and helping in structural remodeling during synaptic plasticity (Calverley and Jones 1990). They are vital in processing large synaptic inputs, and their reduction may impair neuronal transmission and lead to their eventual death (Segal 2010). Singular and combined exposure of MAL and 0.5Gy contributes to neuronal dysfunction by inducing changes in the dendritic morphology, arborization, and dendritic spine density. A search of studies on dendritic morphology analysis post MAL exposure yielded only one other study where MAL (40mg/kg, ip) showed altered dendritic arborization and spine density (Wang et al. 2020), while IR also yielded similar results at 0.1 and 1 Gy doses (Parihar et al. 2015).

### Transcriptomic analysis

Transcriptomic studies can shed light upon crucial changes on a genome-wide level and give valuable insights into the modulation of various cellular processes and altered signaling pathways. So far, there are no reports on the transcriptional regulation of neurotoxicity induced by MAL. MAL exposure altered several biological processes like axonogenesis, neuron differentiation, neuron death, regulation of neuronal synaptic plasticity, forebrain development, regulation of nervous system process, synapse organization, calcium ion homeostasis, axon guidance, regulation of neurotransmitter levels, neural precursor cell proliferation, synapse assembly, regulation of postsynaptic membrane potential, dendrite and dendritic spine development, cognition, negative regulation of neurogenesis, gliogenesis, neuron cellular homeostasis, etc. These changes showed a direct correlation with the histological changes observed in the neuronal morphology, neuronal plasticity, neuroinflammatory response, and neuronal cell death. MAL treatment in rodents has previously been associated with altered dendritic morphology (Campaña et al. 2008), neuroinflammation, apoptotic cell death, and cognitive changes (dos Santos et al. 2016). Several other crucial processes affecting mitochondrial functioning, like ATP metabolic process, mitochondrion organization, mitochondrial respiratory chain complex assembly, oxidative phosphorylation, etc., were also altered. MAL has been shown to induce mitochondrial dysfunction in both *in vivo* and *in vitro* systems (dos Santos et al. 2016; Venkatesan et al. 2017). MAL alters several neurodegenerative pathways like; Pathways of neurodegeneration - multiple diseases that include Huntington’s disease, Alzheimer’s disease, Parkinson’s disease, and Amyotrophic lateral sclerosis. Previous studies linked pesticide exposure to the etiology of neurodegenerative disorders (Sánchez-Santed et al. 2016). Further, MAL was shown to induce several changes in the brain comparable to Alzheimer’s phenotype and was noted to be a good model for studying Alzheimer’s (Venkatesan et al. 2017). Further, pathways affecting related neuronal processes like Glutamatergic synapse, Axon guidance, Spinocerebellar ataxia, and GABAergic synapse were also altered. Among the key signaling pathways affected, the Notch, AMPK, PI3K-Akt, Apelin, Wnt, Calcium, and cAMP signaling pathway were a few of the altered pathways. These pathways have been shown to have a significant influence on the pathogenesis of neurodegenerative diseases (Di Benedetto et al. 2021; Domise and Vingtdeux 2016; Razani et al. 2021; Saito et al. 2019; Ureshino et al. 2019). Interestingly some of these pathways, like the Notch signaling pathway, Wnt signaling pathway, and Apelin signaling pathway, have been reported to be neuroprotective and play a crucial role in modulating neurogenesis and synaptic plasticity (Arrázola et al. 2015; Ho et al. 2020; Zhu et al. 2019). Oxidative phosphorylation was also found to be altered, which is another key pathway involved in neurodegeneration (Chan 2020). Interestingly, the most abundantly altered pathway in our study involved the ribosome and several ribosomal protein-coding genes whose role has so far not been well explored in studies investigating MAL exposure or in studies involving neurodegeneration. Several ribosomal and protein transcription and translation processes, processes related to DNA damage and repair, cell signal activation and transport which included genes like Rpl41, Rpl14, RP19, Rp19-ps6, Rps13, Rps10-ps1, Rpl17, which code for ribosomal proteins involved in large and small ribosomal subunit biogenesis, translation and RNA binding were affected. Studies have shown that proper ribosomal synthesis, continued mRNA translation, and protein production are critical for maintaining the dendritic tree, dynamic proteome remodeling, synapse formation, axon growth, and synaptic plasticity, and impairment can cause disconnection of neuronal circuitries (Slomnicki et al. 2016). Further, ribosomal translational dysregulation is among the important events leading up to the pathogenesis of disorders like Alzheimer’s and Parkinson’s (Ding et al. 2005). Therefore, targeting the ribosomal pathway could be a novel approach to mitigate organophosphate toxicity and hence needs to be further researched. Very few studies have suggested the role of ribosomal proteins in neurodegeneration (Eshraghi et al. 2021; Martin et al. 2014). Most of these have not been reported to be associated with MAL exposure before and hence aid in further understanding MAL’s non-cholinergic mechanism of action.

Singular exposure to radiation altered major biological processes controlling neuronal functioning like axon guidance, regulation of neural precursor cell proliferation, positive regulation of long-term synaptic potentiation, calcium ion homeostasis, neuron apoptotic process, regulation of neuron death, dopamine metabolic process, axonogenesis, synaptic transmission, nerve growth factor signaling pathway, gliogenesis, neurotransmitter loading into synaptic vesicle, dopamine receptor signaling pathway, astrocyte differentiation, neurotransmitter uptake, dopaminergic neuron differentiation, and many more related processes. Interestingly, many dopaminergic processes which play a crucial role among the hallmarks of diseases altered (Michel et al. 2016). Similar to MAL, many cell signaling processes like regulation of MAP kinase activity, cAMP-mediated signaling, regulation of canonical Wnt signaling pathway, ER-nucleus signaling pathway, nerve growth factor signaling pathway, positive regulation of phosphatidylinositol 3-kinase signaling, intrinsic apoptotic signaling pathway, I-kappaB kinase/NF-κB signaling, Notch signaling pathway, hippo signaling, toll-like receptor signaling pathway, etc., were altered. While some of these processes were found to be altered in MAL, some others were unique to IR. The affected processes are involved in the regulation of cellular response to damage through the induction of inflammatory pathways, production of neurotrophins for promoting neurogenesis and neuronal survival, induction of cellular stress response, and repair initiation and neuroprotection. Previously, (Kempf et al.) noted changes in synaptic plasticity, neuronal degeneration, Wnt/β-catenin signaling, and glutamate receptor signaling at 0.3 Gy. Some of the prominent pathways which were enriched IR exposure involved MAPK, PI3K-Akt, Apelin, Calcium, NF-κB, TGF-β, Rap1, Chemokine, cAMP, p53, and Notch signaling pathway, Serotonergic synapse, Axon guidance, Cholinergic synapse, Tyrosine metabolism, and Dopaminergic synapse, etc. Most of the pathways affected were similar to MAL exposure; however, while in MAL, glutamatergic and GABAergic synapses were affected, in low-dose IR exposure, serotonergic and dopaminergic synapse were altered, which are well-known major targets of neurodegeneration (Lowe et al. 2016), has previously reported similar changes in neuronal processes related to neuronal synaptic plasticity and axonal guidance, and altered signaling pathways like cAMP, and ERK/MAPK but also noted glutamate receptor signaling changes and noted similarities in the pathology of Alzheimer’s patients. Some of these processes have been previously reported to be altered in response to low (Kempf et al. 2015; Veeraraghavan et al. 2011). However, many of the signaling processes that have been uniquely altered in our present study have so far been reported to be altered only in high-dose radiation scenarios implying that there is a dearth of information yet to be unearthed in this aspect.

An analysis of the commonly affected biological processes affected between single and combined exposure revealed several processes that regulate neuronal functioning like axon guidance, neuron projection guidance, neural precursor cell proliferation, neuron apoptotic processes, axonogenesis, several protein kinases mediated signaling processes, and DNA damage and repair response. It was noted that the fold enrichment showed an enhanced downregulation of these processes in co-exposure compared to singular exposure to IR and MAL, indicating that MAL and IR interact and reduce the effect and increase neurotoxicity. Similarly, some of the pathways like PI3K-Akt, Ras, and Notch signaling pathways were notable common alterations wherein, compared to the individual exposure, the fold enrichment significantly downregulated in co-exposure, again implying that together, MAL and IR negate the overall effect. Some of the signaling pathways, like the Hippo, FoxO, and JAK-STAT signaling pathway, were found to be uniquely altered only in co-exposure groups. While FOXO, belonging to a family of transcription factors, plays a role in stress response, neuronal signaling, and survival (Santo and Paik 2018), JAK/STAT pathway and Hippo signaling are shown to activate microglia-induced neuroinflammatory response and promote apoptosis (Qin et al. 2016; Sahu and Mondal 2020). Therefore, therapeutic interventions involving inhibition of these pathways could aid in mitigating the neurotoxicity during exposure scenarios. Similarly, GO and pathway enrichment analysis of the set of all the common genes in all three treatment groups revealed that neuronal processes like neurotransmitter metabolic process, neurotransmitter catabolic process, neuron fate specification, neuron remodeling, neuron migration, neural precursor cell proliferation, and many other general processes were altered. Among the significant pathways affected, notable ones were the Cytokine-cytokine receptor interaction, PI3K-Akt signaling pathway, JAK-STAT signaling pathway, and the Neuroactive ligand-receptor interaction pathway. As mentioned previously, dysregulation of both PI3K-Akt and JAK-STAT pathways can hinder normal neuronal functioning, while cytokine activation and release are commonly associated with neurodegenerative disorders (Wang et al. 2015). The PPI network in the co-exposure group revealed changes similar to MAL, where the cluster with highly enriched genes included genes like Rpl28, Gm9493, Rps13, Rps27rt, etc., involved in RNA binding like RNA translation and export, coding for ribonuclear proteins. Similar to IR, the rest of the clusters relatively contained about 3-4 genes and controlled varied processes. Therefore, similar to MAL, the ribosome was the major cellular component affected by co-exposure.

In the present study, though several pathways related to neurons were altered, neither open field test nor novel object recognition for the assessment of exploratory behavior, anxiety, memory, and recognition ability revealed any significant changes suggesting that these molecular alterations may not have manifested in a functional capacity. It is also possible that any associated functional changes would have occurred after a latency period, but since our study only looked at the early brain response to MAL or IR exposure, we may not have seen any significant differences. Further, it is possible that some other behavioral changes like social behavior, fear-induced memory or spatial memory, etc., might have been affected, which was not investigated in our study.

To conclude, single or combined exposure to MAL and IR involves majorly overlapping biological processes and pathways affecting neuronal functions, which could be why they are both implicated in the etiology of neurodegenerative disorders. Our study provides new and detailed insights into the molecular modulation of neurotoxicity associated with MAL and IR exposure and stresses the importance of minimizing exposure to organophosphates like MAL in domestic settings or undergoing routine procedures involving radiation unnecessarily. Further, the improved information can help regulatory authorities revisit and reconsider the previously set safety thresholds and also monitor exposed individuals by looking at various biomarkers, as revealed in our study. Overall, our study suggests that exposure to neurotoxic agents like MAL or IR at a young age, when there are a greater number of developing cells sensitive to xenobiotic damage, could drive the neuronal damage and cell death, thus propelling the occurrence of neurodegenerative complications in the exposed individual at an earlier pace than it usually would occur.

## Acknowledgements

The authors would like to thank the Director, Manipal School of Life Sciences and Manipal Academy of Higher Education (MAHE), for support and infrastructure facilities. This work was supported by The Science and Engineering Research Board Grant No ECR/2017/001239/LS, Government of India. Rekha would like to thank MAHE, Manipal, for the Dr. T.M.A Pai fellowship and KSTePS, Govt of Karnataka, for the scholarship.

## Notes

### Competing Interest Statement

The authors have declared no competing interest.

