## supplementary figure file for "Low-dose exposure to malathion and radiation culminates in the dysregulation of multiple neuronal processes instigating neurotoxicity and activation of neurodegeneration pathways in mice hippocampus"

**Supplementary Figures:**

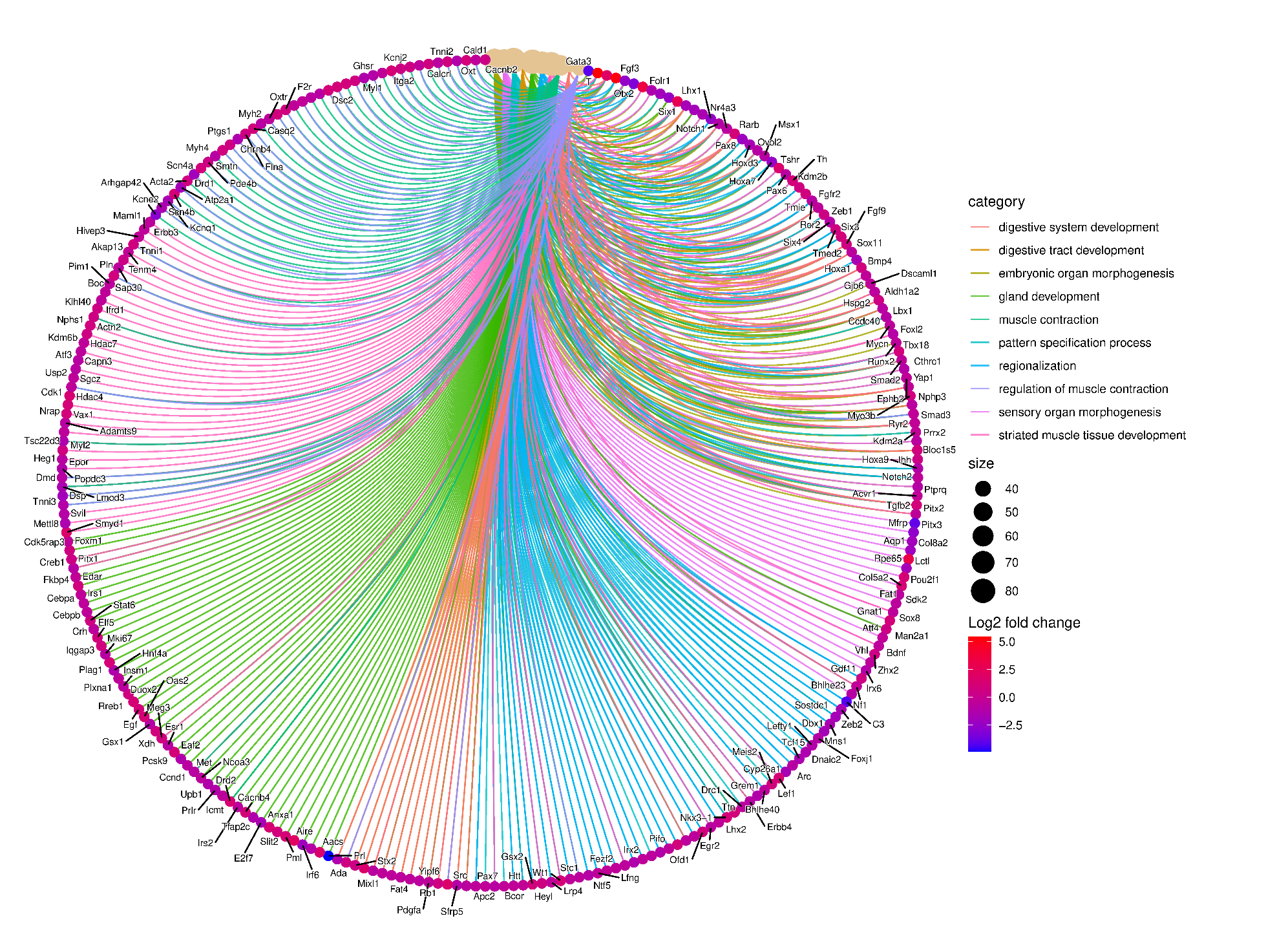

***Fig 1:*** *Cnet plot of top 10 biological processes affected and the respective list of genes involved in radiation group*

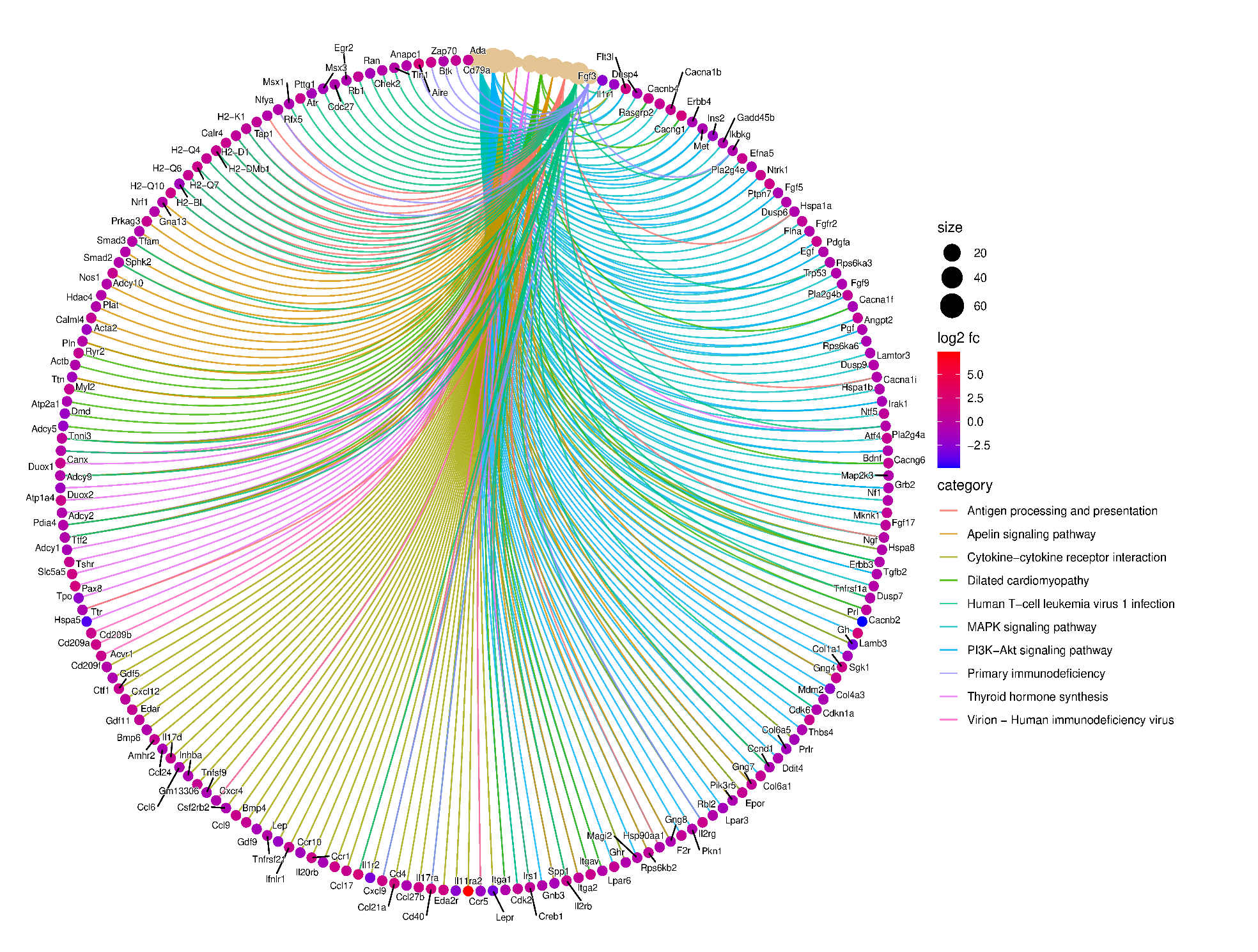

***Fig 2:*** *Cnet plot of top 10 pathways affected and the respective list of genes involved in radiation group*

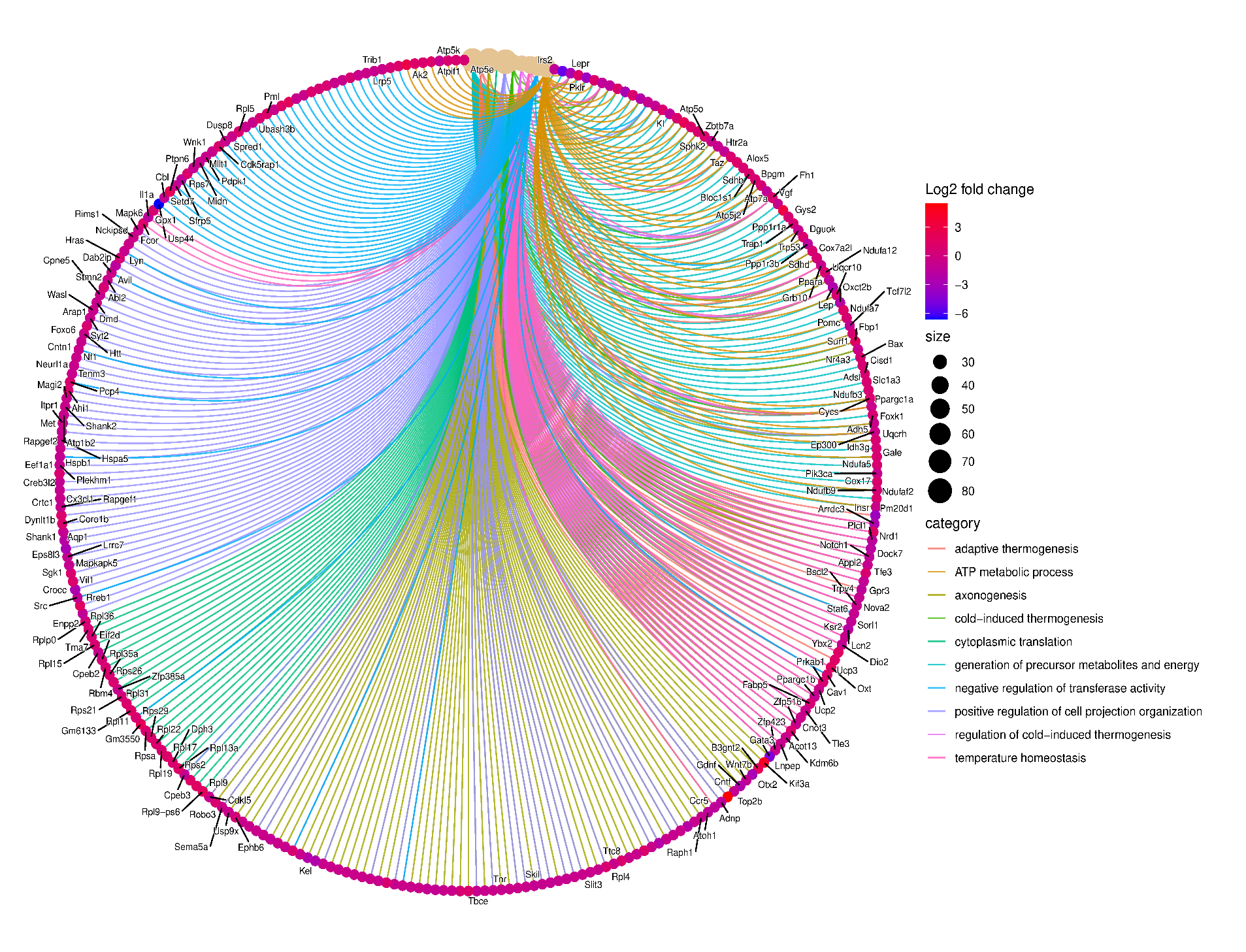

***Fig 3:*** *Cnet plot of top 10 biological processes affected and the respective list of genes involved in malathion group*

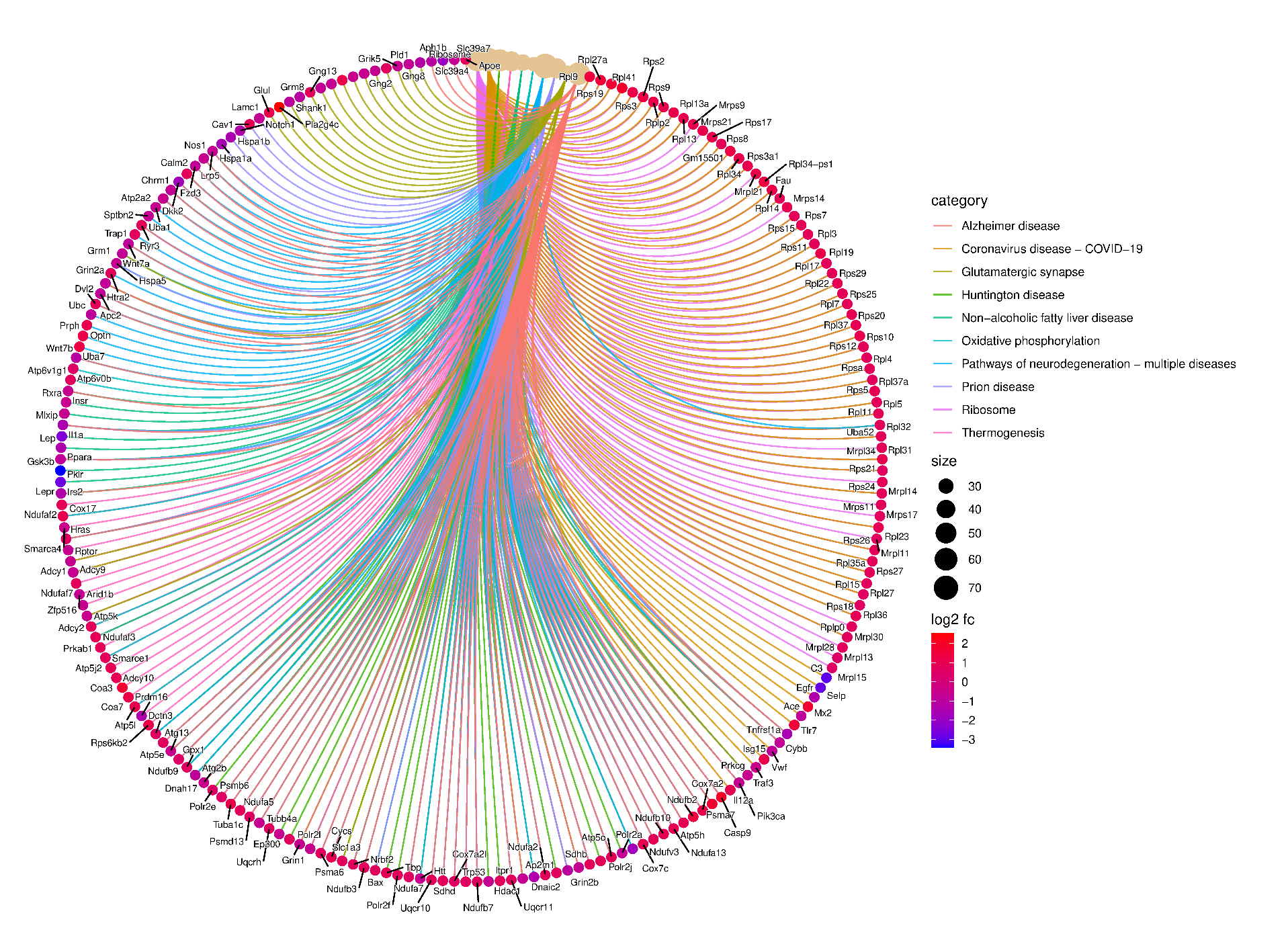

***Fig 4:*** *Cnet plot of top 10 pathways affected and the respective list of genes involved in malathion group*

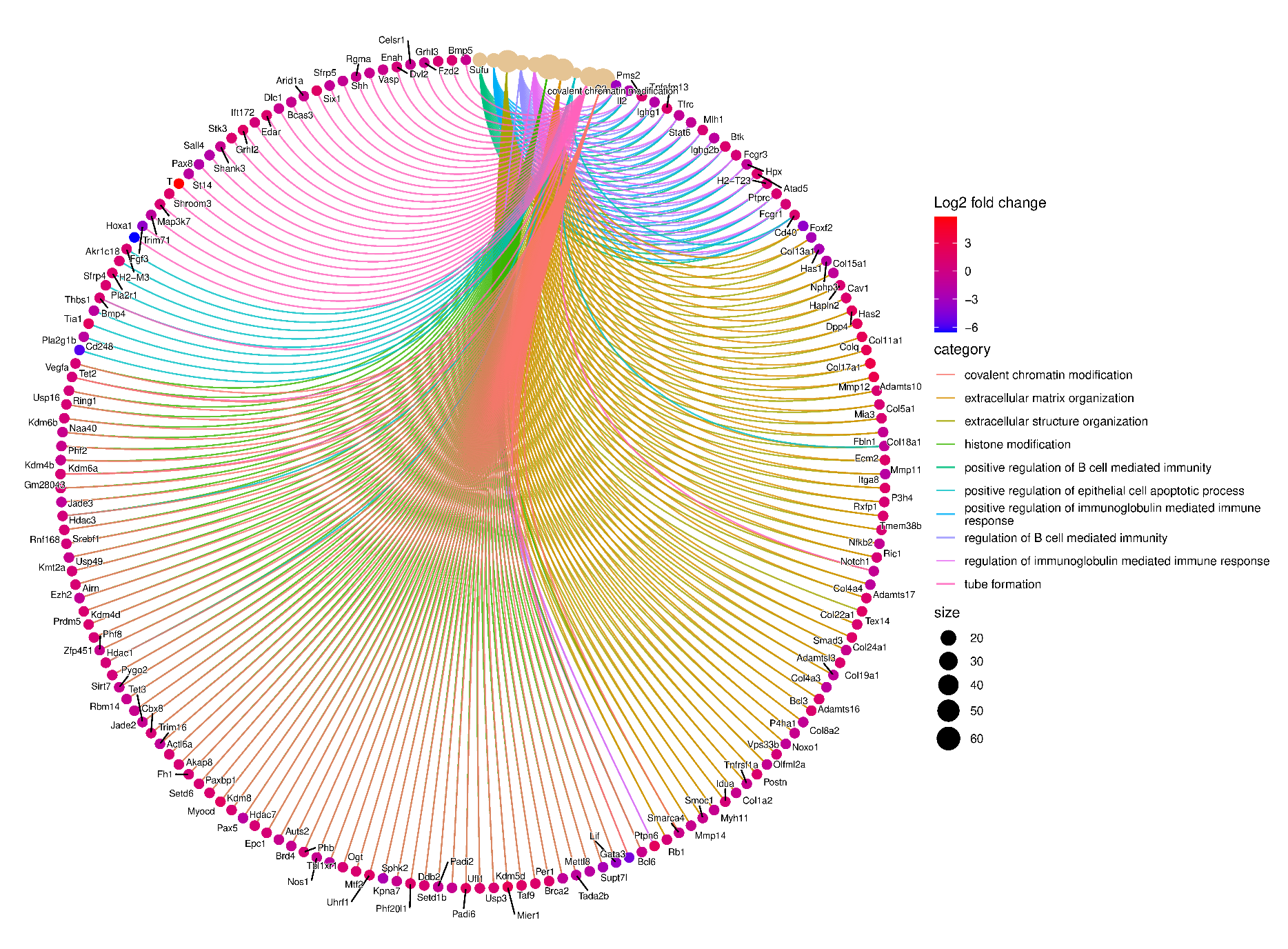

***Fig 5:*** *Cnet plot of top 10 biological processes affected and the respective list of genes involved in co-exposure group*

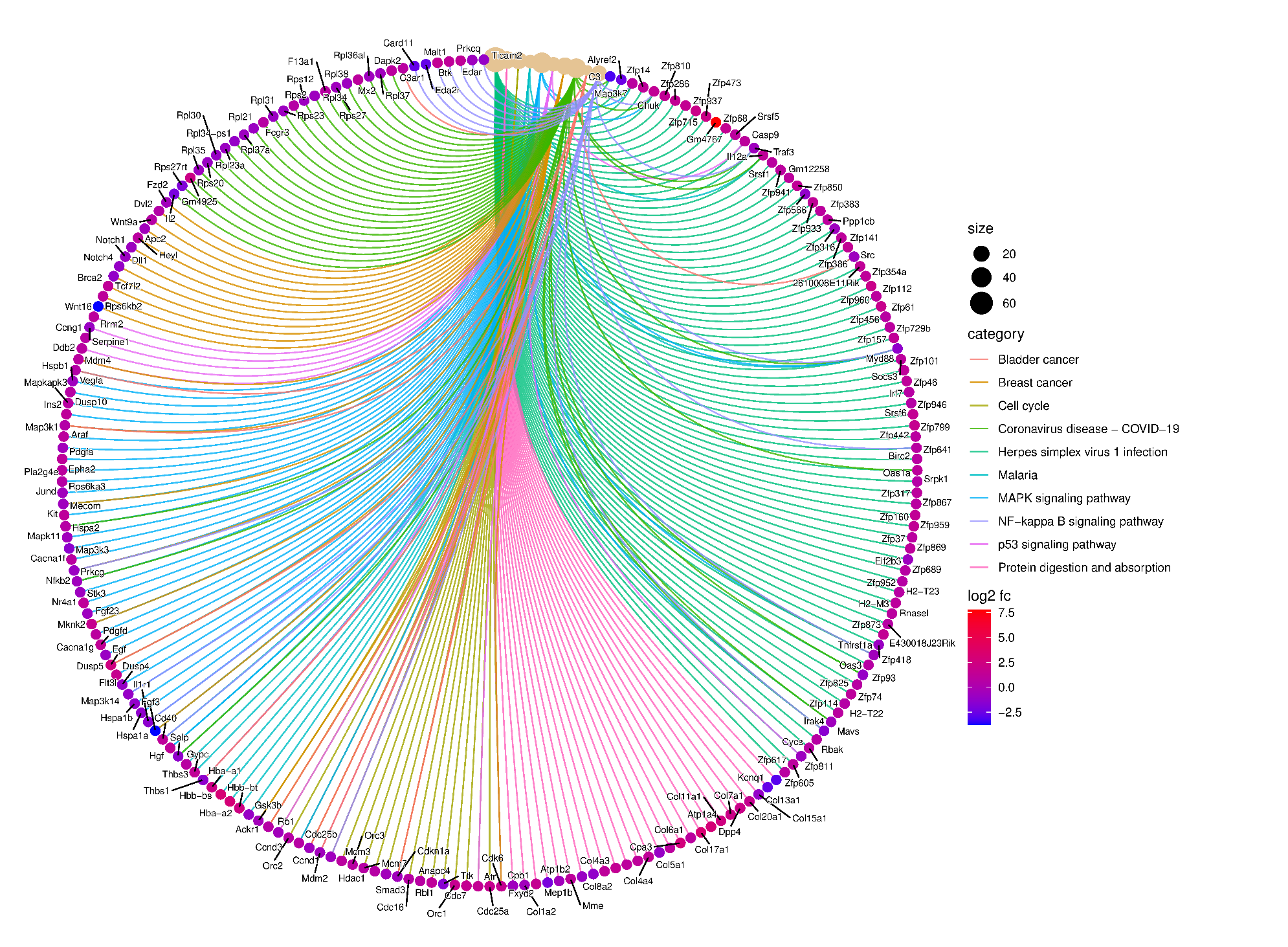

***Fig 6:*** *Cnet plot of top 10 pathways affected and the respective list of genes involved in the co-exposure group*
